# Influence of Pink Pigmented Facultative methylotrophic bacteria (PPFM) as a foliar spraying on the growth and productivity of strawberry *(Fragaria xananassa Duch.)*

**DOI:** 10.1101/2021.12.20.473570

**Authors:** Shadia A. Ismail, A.M. Fafy

**Affiliations:** Potato and Vegetatively Propagated Crops Dep., Hort., Res., Inst., ARC, Giza, Egypt; Botany department, Faculty of women for Arts, Science and Education, Ain Shams University, Cairo, Egypt

**Author notes:** These authors contributed equally to this work. These authors also contributed equally to this work.

**Keywords:** Epiphytes, Methanol, PPFM, Strawberry, Quality and Yield

## Abstract

Pink pigmented facultative methylotrophic bacterium (PPFM) has a favorable impact on plant development and production, it is known as a biostimulator, biofertilizer and biocontroller. Here we investigate the effect of foliar spraying of PPFM, 10% methanol,30% methanol, and their combinations on the growth, fruit quality, and yield of two strawberry cultivars. PPFM was isolated from cotton leaves using imprinting technique. 16S rRNA sequence analysis identified it to be Methylobacterium radiotolerance. Its 16S rRNA sequence were deposited in the Gene Bank under accession number MT644122.1. Two field experiments were conducted during the 2017/2018 and 2018/2019 seasons to investigate the effect of foliar spraying of PPFM and methanol (10 and 30%) on the growth, fruit quality, and yield of two strawberry cultivars. The obtained results showed that, there were no significant differences in the most characteristics between the two cultivars except foliage fresh weight and early yield were higher in cv. Florid, however, Festival cv. recorded higher total yield /plant, anthocyanins and ascorbic acid content in both seasons. Spraying PPFM exhibited the highest values of chlorophyll, fresh weight, total yield and quality. Furthermore, PPFM combined with methanol 10% gave the highest values of leaf area, dry matter %, early yield and some fruit quality. Spraying cv. Florida with PPFM resulted in the best interactions for early yield. However, the best interaction for total yield and most fruit quality features was observed with Festival c.v. and spraying PPFM. It is reasonable to conclude that PPFM is the most effective treatment, increasing strawberry total yield/fed by 28.1 % in the 1st season and 27.91 % in the 2^nd^ season compared to the control.

## Introduction

Strawberry (*Fragaria x ananassa* Duch.) is a highly desirable fruit crop around the world due to its nutrition value, health benefits and export interest. It is an excellent source of natural antioxidants and flavonoids with free radicals scavenging capacity and anticancer properties[1].Strawberry has become one of Egypt’s most important horticultural crops grown of fresh consumption, processing, and exportation. According to Egypt’s Central Administration of Horticulture, Ministry of Agriculture and Land Reclamation, strawberry ranked fifth among exported crops, with a planted area of about 25,000 fed., producing 96,640 tons and exported 36,939 tons in the 2020/2021 season. The above ground phyllo sphere including plant stem and leaves represents a suitable environment for a variety of microorganisms. It emits methanol from stomata as the waste product of pectin metabolism during the early stages of leaf cell elongation. Methylobacterium sp. can utilize methanol as a carbon source for their growth and energy[2].

Methylobacteria, class proteobacteria, consists mainly of a group called Pink pigmented facultative methylotrophic bacteria (PPFM) which are aerobic, Gram-negative bacteria and able to grow on one-carbon compounds such as formate, formaldehyde, and methanol as their sole carbon and energy source[3].Methylobacterium sp. is known as biostimulator, biofertilizer, and biocontrol because it has a positive impact on plant growth and productivity. They enhance plant growth by many mechanism; generating plant growth regulators such as cytokinin and auxins [4] which affect plant growth and various physiological processes, enzymes synthesis such as urease or 1aminocyclopropane-1-carboxylatedeaminase (ACCD) that modulate plant growth [5]. Moreover, they produce phytohormones, provide nutrients to plants and induce systemic resistance in plants against plant pathogens[6].

Methanol is considered a natural product of pectin metabolism in the plant cells. Some of it emits to the atmosphere and the remaining part is converted to formaldehyde, formic acid and finally is converted to CO_2_.The produced CO_2_ affects the CO_2_ assimilation rate in plants and increases secondary metabolites production [7].Increasing CO_2_ concentration inside stomata led to accelerate the rate of photosynthesis and decrease the rate of photorespiration in C plants[8]. Generation of CO_2_ from methanol can also utilized by PPFM [9].

Foliar application of PPFM has a significant effect on the development of many crops including tomato [10], potato [11] and ginger [2], so it can be used for sustainable agriculture. So, the aim of this study is to isolate PPFM and explore its effects as foliar application with methanol, on the growth, yield, and fruit quality of two strawberry cultivars.

## Material and methods

### Isolation and purification of PPFM

A healthy illness symptoms free young cotton leaves collected from the garden of botany department-Faculty of women for Arts, Science and education were used for isolation of PPFM. About 3-4 young leaves were washed using tap water to remove any debris then dried and imprinted on the surface of Ammonium Mineral Salts (AMS) medium supplemented with 0.5% methanol as a sole carbon source (12]. All plates were incubated at 30°C for one week. After a good growth PPFM colony were selected according to its characteristic pink color and further purified on AMS medium and stored for short period on AMS slants at 4°C and for long period as 30% glycerol stock stored at −80°C.

### Identification of PPFM

Methylobacterium isolate used in this study was identified based on the biochemical tests in Bergey’s Manual of Systematic Bacteriology[13]. Molecular identification was carried out using 16SrRNA analysis [14]; Well grown bacterial culture on AMS broth medium was centrifuged at 4000rpm for 10 min. Extraction of DNA was performed using Gen Jet genomic DNA purification Kit (Thermo K0721). Total extracted DNA used as template and amplified by PCR with the aid of oligonucleotide primers;16S rDNA forward primer 8F: (5-AGAGTTTGATCCTGGCTCAG-3), and reverse primer 1520R: (5-TGCGGCTGGATCACCTCCTT-3) were used to obtain nearly 260 bp.

### 16S rDNA sequencing and data analysis

Sequencing analysis was performed on a 255 bp PCR product. The sequence analysis was performed using the ABI 3130 genetic analyzer as well as Big Dye Terminator version 3.1 cycle sequencing kit. The 16S rDNA sequence was aligned and compared with other partial 16S rDNA gene sequences in the GenBank by using the NCBI Basic Local alignment search tools BLAST-n program (http://www.ncbi.nlm.nih.gov/BLAST). Multiple alignments of sequences and nucleotide sequence statistics and variability were performed using CLUSTALW (ver. 1.74) program [15]. Phylogenetic relationship among Egyptian Methylobacterium isolate compared with other international isolates registered in NCBI using Molecular Evolutionary Genetics Analysis (MEGA) software (ver. 4.0) [16]. The accession number MT644122.1 was for the 16S rRNA deposition in the GenBank.

### Inoculum preparation

PPFM isolate was grown on AMS broth medium in an orbital shaker at 120 rpm and 30°C for 10 days. After incubation period bacterial growth was measured using spectrophotometer to be 2 OD at 600 nm[5]. This stock solution was further diluted by mixing 10 ml to 1L of distilled water for plant colonization.

### Field experiment

The investigation was conducted out at the EL-Qanater, Research Station Farm in Qalubia Governorate, Egypt, throughout two seasons, 2017/2018 and 2018/2019 using fresh strawberry transplants (*Fragaria* x *ananassa* Duch.) cv. Festival and Florida. The transplants were obtained from the Strawberry and Non-Traditional Crops Improvement Center of Ain Shams University’s Faculty of Agriculture. Strawberry transplants were planted in the1st and 4th of October in the first and second growing seasons, respectively. Fresh transplants were grown in raised beds 15-20 cm high and 100 cm wide. The transplants were planted 30 cm apart in furrows with drip irrigation. The experimental design was split plot, with two cultivars in the main plot and six randomized foliar spraying treatments in the sub plot, with three replicates and plot area 20 m^2^.

Both cultural activities (irrigation, fertilization, weeding, and pest control) were carried out during both seasons and were performed according to the recommendations of the Egyptian Ministry of Agriculture. The treatments were as following:

1. Distilled water
2. 10% methanol
3. 30% methanol
4. PPFM + 10% methanol
5. PPFM + 30% methanol
6. PPFM

Spraying of the treatments began one month after planting and was repeated six times at three-week intervals and the lower leaf surface was sprayed until wet, as was the upper surface; since the influence of methanol is dependent on a relatively low air temperature in the morning[17].

### Data recorded

#### Vegetative growth

Data were collected from a random sampling of ten plants from the two inner rows of each experimental plot at the start of flowering. The leaf area was measured using the Laser Area Meter CI-202 USA. The fresh weight of the foliage was registered, and it was dried in an oven at 70°C until constant weight to record foliage dry matter percentage. Chlorophyll content was measured as relative values of youngest fourth completely expanded leaf randomly from five plants per plot at four positions on the leaf and then summed them using a handheld chlorophyll meter (SPAD–502, Konica Minolta Sensing, Inc., Japan) [18].The total carbohydrate of crowns was estimated 120 days after planting according to James [19].The potassium content of the leaves were determined using a Flame photometer [20]

#### Yield components

The early yield/plant was calculated using the weights of all harvested fruit during the first four harvests. Total yield/plant and / fed. were measured from the first harvest to the first week of June.

Fruit quality, thirty completely mature fruits were randomly harvested from each treatment in the middle of the growing season (March in both seasons) as subsamples for fruit quality. Fruit firmness were measured with a Chatillon penetrometer. The total soluble solid content (TSS) was measured as described by AOAC [20].

Localization of PPFM strain on the stomata of young plant leaves was confirmed by using scanning electron microscope (JEOL JSM 5200, JEOL Technics Ltd., Japan).

#### Statistical analysis

The MSTAT software was used for the statistical analysis. The data are exposed to an analysis of variance (ANOVA).

## Results and discussion

A PPFM was isolated from the young leaves of cotton plant on AMS medium supplemented with 0.5% methanol as a sole carbon and energy source using leaf impression technique. The isolate was primarily identified as PPFM based on its characteristic pink color[21].Morphologically, this isolate is pink pigmented, aerobic and gram negative bacteria.it was positive catalase, oxidase, urease, Voges-Proskauer (VP), motility and nitrate reduction test. In addition, it was negative for Indole and methyl red (MR) test. The isolate cannot utilize neither starch nor gelatin as carbon source. All these results are in accordance with the phenotypic characteristics of the genus Methylobacterium recorded by Madhaiyan et al [21]

The partial 16S rRNA sequence analysis indicated that the strain belongs to the genus Methylobacteria with 100% similarity with *Methylobacterium radiotolerance*. Its accession number was MT644122.1 as presented in Fig 1.

**Fig 1.**
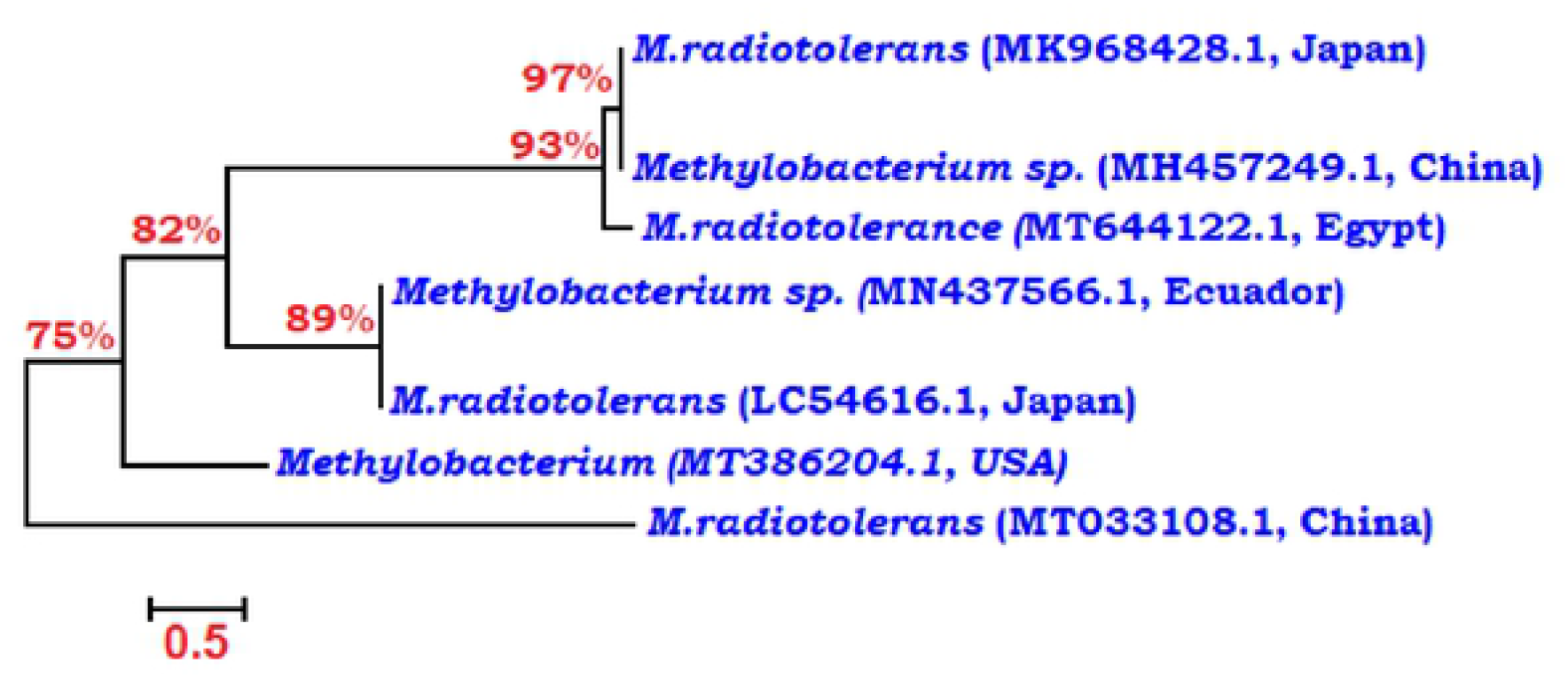
Phylogenetic tree of Methylobacterium isolate based on 16S rRNA sequence analysis. Bootstrap values more than 70% of 1000 Replicates are shown at branch note. Bar, 0.5%.

### Plant height

Table 1 revealed that there was no significant difference between cv. Florida and cv. Festival plants in plant height. Foliar spraying of PPFM bacteria individually or combined with methanol at 10 or 30 % produce the highest plant height in both seasons. However, the lowest values of plant height were recorded for control. Methanol treatment of Florida plants with 30% concentration recorded the highest value of plant heigh. This may be attributed to that, methanol activated the expression of pectin methyl esterase gene that may lead to higher degradation of pectin in the cell wall, producing additional methanol and increased the amount of Ca^2+^ availability, this amount of available Ca^2+^ may be used to promote the growth and the elongation in plant tissues[8]. Moreover, the treatment with PPFM bacteria alone or combined with 10% methanol increased plant height of two cultivars in the two seasons. This might be due to methyl bacteria produce phytohormones such as auxin which promotes plant cell division, extension and stimulate the elongation of plant.

**Table 1.** Effect of Methylobacterium, methanol and their combinations as a foliar spraying on the growth of two strawberry cultivars during 2017/2018 and 2018/ 2019 seasons.

| characters | plant height (cm) |  |  | leaf area (cm<sup>2</sup>) |  |  | Foliage fresh weight (g) |  |  |
| --- | --- | --- | --- | --- | --- | --- | --- | --- | --- |
| cultivars<br>Treatment | Florida | Festival | Mean | Florida | Festival | Mean | Florida | Festival | Mean |
| **2017-2018** |  |  |  |  |  |  |  |  |  |
| **Control** | 20.19<br>f | 19.27<br>f | 19.7<br>C | 783.3<br>ef | 673.6<br>g | 728.5<br>D | 53.53<br>f | 34.68<br>l | 44.1<br>D |
| **10%methanol** | 22.27<br>e | 23.67<br>cd | 22.9<br>B | 976.6<br>b | 848.7<br>d | 912.7<br>B | 60.89<br>e | 42.05<br>j | 51.7<br>C |
| **30% methanol** | 24.83<br>a | 23.03<br>de | 23.9<br>3 A | 862.3<br>cd | 785.1<br>e | 823.7<br>C | 63.18<br>d | 40.43<br>k | 51.8<br>1 C |
| **PPFM+**<br>**10%methanol** | 23.73<br>bcd | 24.80<br>a | 24.3<br>A | 1031.<br>a | 886.6<br>c | 958.8<br>A | 65.71<br>b | 46.12<br>h | 55.9<br>AB |
| **PPFM+30**<br>**%methanol** | 24.20<br>abc | 22.30<br>e | 23.3<br>B | 871.7<br>cd | 758.4<br>f | 815.0<br>C | 64.95<br>c | 45.34<br>i | 55.2<br>B |
| **PPFM** | 23.97<br>abc | 24.63<br>ab | 24.3<br>0 A | 987.1<br>b | 857.1<br>d | 922.1<br>AB | 66.62<br>a | 46.98<br>g | 56.8<br>A |
| **Mean** | 23.2<br>A | 22.95<br>A |  | 918.7<br>A | 801.6<br>A |  | 62.48<br>A | 42.60<br>B |  |
| **2018-2019** |  |  |  |  |  |  |  |  |  |
| **Control** | 20.63<br>f | 22.17<br>e | 21.4<br>0 C | 797.4<br>d | 686.9<br>e | 742.2<br>D | 58.16<br>e | 37.51<br>k | 47.8<br>4 E |
| **10%methanol** | 24.70<br>d | 24.93<br>d | 24.8<br>2 B | 998.9<br>b | 864.1<br>c | 931.5<br>BC | 65.84<br>d | 45.53<br>i | 55.6<br>8 C |
| **30%methanol** | 26.63<br>ab | 25.83<br>c | 26.2<br>3 A | 876.2<br>c | 866.5<br>c | 871.4<br>C | 66.83<br>c | 42.78<br>j | 54.8<br>1 D |
| **PPFM+**<br>**10%methanol** | 26.90<br>a | 26.00<br>bc | 26.4<br>5 A | 1121<br>a | 991.2<br>b | 1056.<br>1 A | 71.56<br>a | 49.52<br>h | 60.5<br>4 B |
| **PPFM+**<br>**30%methanol** | 24.97<br>d | 25.83<br>c | 25.4<br>AB | 1005<br>b | 875.6<br>c | 940.2<br>BC | 68.92<br>b | 51.21<br>g | 60.1<br>B |
| **PPFM** | 26.10<br>bc | 26.37<br>abc | 26.2<br>3 A | 1049<br>b | 1003<br>b | 1026<br>AB | 71.93<br>a | 52.11f | 62.0<br>2 A |
| **Mean** | 25.07<br>A | 25.31<br>A |  | 974.6<br>A | 881.2<br>A |  | 67.21<br>A | 46.44<br>B |  |

Values within the column or rows followed by the same capital or small letter/s do not significantly differ from each other according to Duncan's multiple range test at 5% level.

### Leaf area

Data in Table 1 show that there was no significant difference of leaf area between cv. Florida and cv. Festival. Spraying strawberry plants with PPFM combined with 10% methanol gave the highest leaf area in both growing seasons. Also, spraying strawberry cv. Florida with PPFM combined with 10% methanol revealed the maximum interaction of leaf area in both seasons. These findings may be attributed to methylotrophic bacteria, which use the gaseous methanol released by plants through the stomata as a carbon and energy supply while also promoting the growth of their host through the release of growth promoting substances such as auxin and cytokinin, which play an important role in regulating plant growth and development by promoting cell division proliferation, growth (expansion, elongation and differentiation)[22]. Methanol foliar spraying slowed leaf senescence in the plant by increasing photosynthetic active time and leaf area span[23].Also, it plays an important function in controlling cell expansion by activating cell wall synthesis [24,25].

### Foliage fresh weight

Florida c.v. plants have the highest foliage fresh weight value compared with Festival c.v. Strawberry plants treated with PPFM have the highest foliage fresh weight values in both seasons followed by methanol treatments in the second season Table 1. Our results are consistent with those of Nadali et al. [4]. Florida plants treated with PPFM or mixed with 10% methanol resulted in the maximum interaction values of foliage fresh weight in the first season. These findings are in harmony with those obtained by Abbasian et al [26].Adding methanol to Methylobacteria might have positive effect for increasing carbon dioxide in the leaves, causing an increase in photosynthetic production in the leaves, resulting in a delay in leaf senescence by preventing plant ethylene levels from being growth inhibitory[23].

### Chlorophyll content

Florida c.v. and cv. Festival had similar chlorophyll content measurements during both seasons (Table 2). Spraying strawberry plants with PPFM resulted in the highest chlorophyll content for the two seasons. Methylobacterium able to improve photosynthetic activity by increasing stomatal count and chlorophyll content in rice plants[27]. Spraying methanol has a beneficial impact when combined with Methylobacteria [24]. Florida c.v. plants sprayed with PPFM had the highest interaction chlorophyll values in both seasons followed by Festival c.v. plants with PPFM in the second season.

**Table 2.** Effect of methylobacterium, methanol and their combinations as a foliar spraying on the chlorophyll content, foliage dry matter percentage and carbohydrates of two strawberry cultivars during 2017/2018 and 2018/ 2019 seasons.

| Characters | Chlorophyll content (SPAD) |  |  | Foliage dry matter (%) |  |  | Carbohydrates (mg/g dry weight) |  |  |
| --- | --- | --- | --- | --- | --- | --- | --- | --- | --- |
| cultivars<br>Treatments | Florida | Festival | Mean | Florida | Festival | Mean | Florida | Festival | Mean |
| **2017-2018** |  |  |  |  |  |  |  |  |  |
| **Control** | 46.66<br>i | 48.67<br>h | 47.66<br>D | 20.72<br>f | 18.91<br>g | 19.81<br>D | 4.670<br>f | 4.787<br>e | 4.728<br>D |
| **10%methanol** | 49.77<br>g | 51.57<br>cde | 50.67<br>C | 24.80<br>c | 22.89<br>e | 23.84<br>BC | 5.653<br>b | 5.557<br>c | 5.605<br>B |
| **30%methanol** | 50.70<br>f | 50.57<br>f | 50.63<br>C | 23.92<br>d | 22.54<br>e | 23.23<br>C | 5.446<br>d | 5.542<br>c | 5.494<br>C |
| **PPFM+**<br>**10%methanol** | 51.93<br>bcd | 52.22<br>b | 52.08<br>AB | 25.79<br>b | 25.76<br>b | 25.78<br>A | 5.694<br>b | 5.935<br>a | 5.815<br>A |
| **PPFM+**<br>**30%methanol** | 51.33<br>de | 51.07<br>ef | 51.20<br>BC | 25.10<br>c | 23.90<br>d | 24.50<br>B | 5.652<br>b | 5.559<br>c | 5.606<br>B |
| **PPFM** | 53.00<br>a | 52.13<br>bc | 52.57<br>A | 26.45<br>a | 24.26<br>d | 25.35<br>A | 5.672<br>b | 5.879<br>a | 5.777<br>A |
| **Mean** | 50.57<br>A | 51.04<br>A |  | 24.46<br>A | 23.04<br>A |  | 5.464<br>A | 5.544<br>A |  |
| **2018-2019** |  |  |  |  |  |  |  |  |  |
| **Control** | 50.80<br>g | 51.10<br>g | 50.95<br>E | 21.79<br>f | 21.49<br>f | 21.64<br>C | 4.379<br>g | 4.583<br>f | 4.481<br>D |
| **10%methanol** | 51.13<br>g | 53.20<br>cd | 52.17<br>CD | 26.30<br>c | 26.04<br>cd | 26.17<br>B | 5.603<br>d | 5.466<br>e | 5.534<br>C |
| **30%methanol** | 52.37<br>e | 51.67<br>f | 52.02<br>D | 25.46<br>de | 25.94<br>cd | 25.70<br>B | 5.823<br>c | 5.589<br>d | 5.706<br>B |
| **PPFM+**<br>**10%methanol** | 53.17<br>cd | 53.50<br>c | 53.33<br>B | 27.70<br>a | 27.30<br>ab | 27.50<br>A | 6.452<br>b | 5.831<br>c | 6.142<br>A |
| **PPFM+30%**<br>**methanol** | 52.93<br>d | 52.80<br>de | 52.87<br>BC | 26.66<br>bc | 24.83<br>e | 25.75<br>B | 6.516<br>b | 5.857<br>c | 6.187<br>A |
| **PPFM** | 55.87<br>a | 54.57<br>b | 55.22<br>A | 27.91<br>a | 27.43<br>ab | 27.67<br>A | 5.619<br>d | 6.746<br>a | 6.183<br>A |
| **Mean** | 52.71<br>A | 52.81<br>A |  | 25.97<br>A | 25.50<br>A |  | 5.73<br>A | 5.68<br>A |  |

Values within the column or rows followed by the same capital or small letter/s do not significantly differ from each other according to Duncan's multiple range test at 5% level.

### Foliage dry matter %

Results in Table 2 show that there was no substantial difference in foliage dry matter percentage between cv. Florida and cv. Festival. Spraying strawberry plants with PPFM or PPFM mixed with10% methanol provided the highest values of foliage dry matter percentage. Furthermore, either PPFM combined with 30% methanol or methanol treatments showed a higher significant increase compared with control in both seasons. In this regard, increasing in dry matter and its translocation is linked to hormonal balance, and PPFM synthesizes a range of auxins and cytokinin that are used by host plants for development[28]. For Florida plants, PPFM resulted in the highest interaction values of foliage dry matter in the first season followed by PPFM mixed with 10 % methanol for Festival plants. These results are consistent with those obtained by[29,30,31) in strawberry and lavender plants, respectively.

### Crown carbohydrates content

Data presented in Table 2 show that there were no significant differences in carbohydrates content between the two strawberry cultivars and PPFM treatment either individually or combined with 10% methanol resulted in the highest values of carbohydrates content in the two growing seasons. This result is due to the photosynthesis increase by PPFM and photorespiration reduction by raising the concentration of CO surrounding and within the stomata that increase carbohydrate synthesis[8]. Foliar application of methanol at 10 % and 30% had a positive impact on carbohydrate content as compared to the control. PPFM provided the highest interaction values of carbohydrates content of Florida c.v. in the two seasons. [32] found that methanol gave the highest increase carbohydrates of Flame seedless grapevines.

### Early yield

Florida plants gave more early yield compared with cv. Festival plants in both growing seasons (Table 3). This results are in accordance with that recorded [33]

**Table 3.** Effect of methylobacterium, methanol and their combinations as a foliar spraying on the yield of two strawberry cultivars during 2017/2018 and 2018/ 2019 seasons.

| Characters<br>cultivars<br>Treatments | Early yield(g/plant) |  |  | Total yield(g/plant) |  |  | Total yield (ton/fed) |  |  |
| --- | --- | --- | --- | --- | --- | --- | --- | --- | --- |
|  | Florida | Festival | Mean | Florida | Festival | Mean | Florida | Festival | Mean |
| **2017-2018** |  |  |  |  |  |  |  |  |  |
| **Control** | 190.7<br>d | 130.9<br>e | 160.8<br>C | 426.2<br>l | 473.8<br>k | 450.0<br>E | 16.98<br>l | 18.95<br>k | 17.97<br>E |
| **10%methanol** | 224.8<br>c | 195.6<br>d | 210.2<br>B | 510.5<br>j | 559.4<br>e | 534.9<br>D | 20.42<br>j | 22.31<br>e | 21.36<br>D |
| **30%methanol** | 235.2<br>bc | 227.7<br>c | 231.4<br>AB | 515.2<br>i | 569.1<br>c | 542.2<br>C | 20.60<br>i | 22.76<br>c | 21.68<br>C |
| **PPFM+**<br>**10%methanol** | 239.9<br>abc | 231.9<br>bc | 235.9<br>AB | 548.7<br>g | 594.8<br>b | 571.7<br>B | 21.88<br>g | 23.79<br>b | 22.84<br>B |
| **PPFM+**<br>**30%methanol** | 251.9<br>ab | 240.9<br>abc | 246.4<br>AB | 528.9<br>h | 561.9<br>d | 545.4<br>C | 21.12<br>h | 22.48<br>d | 21.80<br>C |
| **PPFM** | 257.9<br>a | 244.0<br>abc | 251.0<br>A | 552.4<br>f | 598.9<br>a | 575.6<br>A | 22.08<br>f | 23.96<br>a | 23.02<br>A |
| **Mean** | 233.4<br>A | 211.8<br>B |  | 513.6<br>B | 559.6<br>A |  | 20.51<br>A | 22.37<br>A |  |
| **2018-2019** |  |  |  |  |  |  |  |  |  |
| **Control** | 192.6<br>e | 142.2<br>h | 167.4<br>D | 452.6<br>k | 488.1<br>j | 470.4<br>F | 18.10<br>k | 19.52<br>j | 18.81<br>F |
| **10%methanol** | 231.4<br>c | 169.7<br>g | 200.5<br>C | 538.3<br>h | 571.8<br>f | 555<br>E | 21.52<br>h | 22.87<br>f | 22.19<br>E |
| **30%methanol** | 235.2<br>bc | 174.1<br>g | 204.7<br>BC | 535.6<br>i | 582.1<br>d | 558.8<br>D | 21.43<br>I | 23.27<br>d | 22.35<br>D |
| **PPFM+**<br>**10%methanol** | 255.3<br>a | 192.5<br>e | 223.9<br>A | 570.6<br>f | 620.8<br>b | 598.5<br>B | 22.85<br>f | 24.99<br>b | 23.92<br>B |
| **PPFM+**<br>**30%methanol** | 239.1<br>b | 184.2<br>f | 211.7<br>B | 546.9<br>g | 588.3<br>c | 567.6<br>C | 21.88<br>g | 23.50<br>c | 22.69<br>C |
| **PPFM** | 259.3<br>a | 200.7<br>d | 230.0<br>A | 578.5<br>e | 626.8<br>a | 602.7<br>A | 23.05<br>e | 25.07<br>a | 24.06<br>A |
| **Mean** | 235.5<br>A | 177.2<br>B |  | 537.4<br>B | 580.3<br>A |  | 21.47<br>B | 23.21<br>A |  |

Values within the column or rows followed by the same capital or small letter/s do not significantly differ from each other according to Duncan's multiple range test at 5% level.

The highest early yield was achieved by PPFM alone or combined with 10% methanol in both seasons. These results could be attributed to Methylbacteria’s ability to prevent early leaf senescence, which increased leaf area and photosynthetic pigments and the role of cytokinin in promoting cell division and fruit set. Spraying Florida plants with PPFM alone produced the highest values of early yield with no significant differences with 30% methanol and mixed treatments in the first season. For the second season, either PPFM alone or combined with 10% methanol were the highest. When methylobacteria were combined with methanol, the amount of methanol consumed increased, which had a positive effect on phytohormone synthesis and promoted the growth of their host via the release of metabolites[3]. Similarly, methanol has been linked to improved growth and yield of marigold plants [34].

### Total yield

Festival plants recorded higher total yield/plant than cv. Florida plants over the two growing seasons Table 3. Furthermore, there was no significant difference in total yield fed/plot between the two cultivars in the first season, however, in the second season, Festival plants recorded a higher average. Treated with PPFM resulted in the highest total yield /plant and fed/plot in the two tested seasons, followed by PPFM mixed with 10% methanol. Festival c.v. treated with PPFM led to a significant increase interaction value of total yield/plant and fed. in both tested seasons. This increase could be attributed to PPFM, which produces phytohormones, provides nutrients, and induces defense responses against plant pathogens. Also, the availability of methanol excreted from the growing plant, encouraging the PPFM to colonize as recorded in potato plants by Ardanov et al[35].

### Fruit firmness

Data in Table 4 indicate that there was no substantial difference in firmness between the two cultivars in the two seasons and PPFM treatment achieved the highest fruit firmness, with no significant differences with either 10% methanol or PPFM combined with 30% methanol in both seasons. This may be due to a rise in cell wall rigidity which resulted in methanol which decreased the activity of antioxidant enzymes and increased [8]. Also, PPFM treatment of Festival plants resulted in the maximum interaction values with no noticeable difference between 10% methanol and PPFM combined with 30% methanol in both studied seasons as recorded by Abo-Sedera et al[29].

**Table 4.** Effect of methyl bacterium, methanol, and their combinations as a foliar spraying on the Firmness, TSS and acidity of two strawberry cultivar Fruits during 2017/2018 and 2018/ 2019 seasons.

| characters | Firmness(kg/cm) |  |  | T.S.S(%) |  |  | Acidity(mg/100g f.w) |  |  |
| --- | --- | --- | --- | --- | --- | --- | --- | --- | --- |
| cultivars<br>Treatments | Florida | Festival | Mean | Florida | Festival | Mean | Florida | Festival | Mean |
| 2017-2018 |  |  |  |  |  |  |  |  |  |
| Control | 228.3<br>f | 260<br>e | 244.2<br>C | 9.200<br>f | 8.733<br>g | 8.967<br>C | 0.8937<br>b | 0.8853<br>b | 0.8895<br>A |
| 10%methanol | 281.7<br>cd | 345<br>ab | 313.3<br>A | 10.80<br>c | 10.27<br>d | 10.53<br>B | 0.8237<br>c | 0.9113<br>ab | 0.8675<br>A |
| 30%methanol | 275<br>de | 288.3<br>cd | 281.7<br>B | 11.60<br>a | 9.800<br>e | 10.70<br>B | 0.93<br>a | 0.8157<br>cd | 0.8728<br>A |
| PPFM+<br>10%methanol | 328.3<br>b | 293.3<br>cd | 310.8<br>AB | 10.60<br>c | 10.00<br>de | 10.30<br>B | 0.8157<br>cd | 0.7950<br>d | 0.8053<br>B |
| PPFM+<br>30%methanol | 280.0<br>d | 343.3<br>ab | 311.7<br>AB | 11.67<br>a | 11.27<br>b | 11.47<br>A | 0.6347<br>f | 0.9320<br>a | 0.7833<br>B |
| PPFM | 300.0<br>c | 361.7<br>a | 330.8<br>A | 11.60<br>a | 11.60<br>a | 11.60<br>A | 0.6670<br>e | 0.6457<br>ef | 0.6563<br>C |
| Mean | 282.2<br>A | 315.3<br>A |  | 10.91<br>A | 10.28<br>A |  | 0.7941<br>A | 0.8308<br>A |  |
| 2018-2019 |  |  |  |  |  |  |  |  |  |
| Control | 253.3<br>f | 271.7<br>e | 262.5<br>C | 9.867<br>f | 9.867<br>f | 9.867<br>D | 0.9667<br>b | 1.021<br>a | 0.9937<br>A |
| 10%methanol | 286.7<br>d | 368.3<br>a | 327.5<br>AB | 11.13<br>de | 11.00<br>de | 11.07<br>C | 0.9397<br>bc | 0.9343<br>c | 0.9370<br>B |
| 30%methanol | 320.0<br>bc | 315.0<br>bc | 317.5<br>B | 12.20<br>a | 11.33<br>cd | 11.77<br>AB | 0.8747<br>d | 0.8450<br>e | 0.8598<br>C |
| PPFM+<br>10%methanol | 318.3<br>bc | 306.7<br>c | 312.5<br>B | 11.73<br>b | 10.87<br>e | 11.30<br>BC | 0.7667<br>f | 0.7563<br>fg | 0.7615<br>D |
| PPFM+<br>30%methanol | 310.0<br>c | 370.0<br>a | 345.0<br>A | 12.30<br>a | 11.73<br>b | 12.02<br>A | 0.6840<br>h | 0.8293<br>e | 0.7567<br>D |
| PPFM | 324.0<br>b | 380.0<br>a | 347.0<br>A | 12.27<br>a | 11.60<br>bc | 11.93<br>A | 0.7280<br>g | 0.7653<br>f | 0.7467<br>D |
| Mean | 302.1<br>A | 335.3<br>A |  | 11.58<br>A | 11.07<br>A |  | 0.827<br>A | 0.859<br>A |  |

Values within the column or rows followed by the same capital or small letter/s do not significantly differ from each other according to Duncan's multiple range test at 5% level.

### T.S.S fruit %

Data in Table 4 indicate that there was no significant difference in total soluble solids between the two cultivars. PPFM,30% methanol and combined treatment of PPFM and 30% methanol provided the highest total soluble solids compared to the control in the two seasons. This result may be related to the fact that methanol is easily metabolized in plant stomata and utilized in photosynthesis, resulting in enhanced sugar production [4]. Also, these treatments provided the highest interaction values for Florida plants in the two seasons. For cv. Festival, PPFM alone or combined with 30%methanol gave the highest values.

### Fruit acidity

There was no significant difference in acidity content between cv. Florida and cv. Festival fruits in the two growing seasons Table 4. In the first season, PPFM treatment provided the lowest value of acidity content in contrast to PPFM and PPFM mixed with methanol in the second season. Ardanov et al [36] and Zabetakis [37] recorded that, Strawberry plants associated with methylobacteria may improve the biosynthesis of certain compounds that affect strawberry fruit quality. For Florida plants PPFM combined with 30 % methanol had the lowest interaction values of acidity in the first season.

### Anthocyanins and ascorbic acid fruit content

Festival fruits had a higher anthocyanin and ascorbic acid content than Florida fruits in both tested seasons Table 5. Our findings are consistent with those of Abo-Sedera et al[29].Moreover PPFM produced the highest anthocyanins and ascorbic acid values during the two studied seasons. Ramadan and Omran [38] recorded that the methanol application improved total anthocyanins in berry skins. Cruz-Rus et al[39]attributed the rise in ascorbic acid to increased synthesis of the six-carbon sugar derivative that is essential for its biosynthesis. In addition, Festival plant treated with PPFM in both seasons resulted in the highest interaction of anthocyanin content followed by PPFM combined with 10 %methanol compared with control. This finding was in accordance with Rahman et al[40] and Valizadeh-Kamran et al [31]. Also, the highest interaction values of ascorbic acid content for Festival c.v. was obtained by PPFM that is may be attributed to its ability to produce stimulators such as auxin, cytokinin, and vitamin B12, which could increase the biosynthesis pathway of ascorbic acid production [28].

**Table 5.** Effect of methylobacterium, methanol and their combinations as a foliar spraying on anthocyanins, ascorbic acid and K content of two strawberry cultivar Fruits during 2017/2018 and 2018/ 2019 seasons.

| Characters | Total anthocyanins<br>mg/100g f. w) |  |  | Ascorbic acid<br>mg/100g (f. w) |  |  | K% |  |  |
| --- | --- | --- | --- | --- | --- | --- | --- | --- | --- |
| cultivars<br>Treatments | Florida | Festival | Mean | Florida | Festival | Mean | Florida | Festival | Mean |
| **2017-2018** |  |  |  |  |  |  |  |  |  |
| **Control** | 65.32<br>i | 90.93<br>h | 78.12<br>D | 39.69<br>h | 53.61<br>d | 46.65<br>D | 1.741<br>e | 1.384<br>g | 1.563<br>C |
| **10%methanol** | 92.19<br>h | 127.5<br>c | 109.8<br>C | 44.73<br>g | 60.97<br>b | 52.85<br>C | 1.861<br>d | 1.500<br>f | 1.680<br>B |
| **30%methanol** | 106.7<br>e | 123.6<br>d | 115.2<br>B | 47.06<br>f | 60.86<br>b | 53.96<br>C | 1.881<br>d | 1.560<br>f | 1.721<br>B |
| **PPFM+**<br>**10%methanol** | 98.88<br>g | 129.9<br>b | 114.4<br>B | 52.02<br>e | 63.21<br>a | 57.61<br>B | 2.058<br>b | 1.759<br>e | 1.908<br>A |
| **PPFM+**<br>**30%methanol** | 98.52<br>g | 129.0<br>b | 113.7<br>B | 52.76<br>de | 63.07<br>a | 57.92<br>AB | 1.954<br>c | 1.530<br>f | 1.742<br>B |
| **PPFM** | 103.2<br>f | 133.5<br>a | 118.4<br>A | 55.11<br>c | 63.88<br>a | 59.49<br>A | 2.199<br>a | 1.760<br>e | 1.979<br>A |
| **Mean** | 94.14<br>B | 122.4<br>A |  | 48.56<br>B | 60.93<br>A |  | 1.949<br>A | 1.582<br>A |  |
| **2018-2019** |  |  |  |  |  |  |  |  |  |
| **Control** | 73.55<br>l | 94.29<br>k | 83.92<br>E | 42.07<br>i | 56.01<br>d | 49.04<br>D | 1.828<br>h | 1.424<br>i | 1.626<br>D |
| **10%methanol** | 98.73<br>j | 129.7<br>e | 114.2<br>D | 47.48<br>h | 61.92<br>c | 54.70<br>C | 1.948<br>fg | 1.973<br>f | 1.961<br>C |
| **30%methanol** | 102.3<br>i | 139.6<br>c | 120.9<br>C | 49.19<br>g | 62.47<br>c | 55.83<br>C | 2.167<br>e | 1.910<br>g | 2.039<br>C |
| **PPFM+**<br>**10methanol** | 109.8<br>h | 144.2<br>a | 127.0<br>A | 53.92<br>e | 64.85<br>a | 59.38<br>AB | 2.450<br>b | 2.405<br>b | 2.427<br>A |
| **PPFM+**<br>**30%methanol** | 114.8<br>f | 134.7<br>d | 124.8<br>B | 52.72<br>f | 63.69<br>b | 58.20<br>B | 2.242<br>d | 2.178<br>e | 2.210<br>B |
| **PPFM** | 111.7<br>g | 141.3<br>b | 126.5<br>A | 55.80<br>d | 64.67<br>a | 60.24<br>A | 2.507<br>a | 2.295<br>c | 2.401<br>A |
| **Mean** | 101.8<br>B | 130.6<br>A |  | 50.20<br>B | 62.27<br>A |  | 2.190<br>A | 2.031<br>A |  |

Values within the column or rows followed by the same capital or small letter/s do not significantly differ from each other according to Duncan, s multiple range test at 5% level.

### K leaf content

Table 5 show that there were no significant differences in leaf K leaves content between the two strawberry cultivars and PPFM greatly improved leaves contents of K during the two growing seasons. Spraying of PPFM or combining it with 10% methanol greatly increased K% in the two tested seasons. In this regard, methylobacterium sp., is a biostimulator and a biofertilizer, and they directly influence plant growth by supplying nutrients to the plants [6]. The highest interaction values of potassium content were recorded by PPFM treated Florida plants in both studied seasons.

### SEM morphological studies

Fig 2 illustrates SEM images of untreated plants and foliar applications of PPFM, either alone or in combination with methanol rates, on strawberry leaves. Fig 2B and 2C show that PPFM can exist in the presence of methanol and PPFM is found in the pore and is associated with stomata. The strawberry leaf became hypersensitive after being treated with methanol (Figs 2B and 2C). When PPFM was combined with Methanol (Figs 2D and 2E), the harmful effects of methanol on leaf tissues were reduced. Also, when PPFM was used alone, the regular structure of strawberry epidermal cells was maintained and most of the investigated characters showed beneficial effects (Fig 2F).

**Fig 2.**
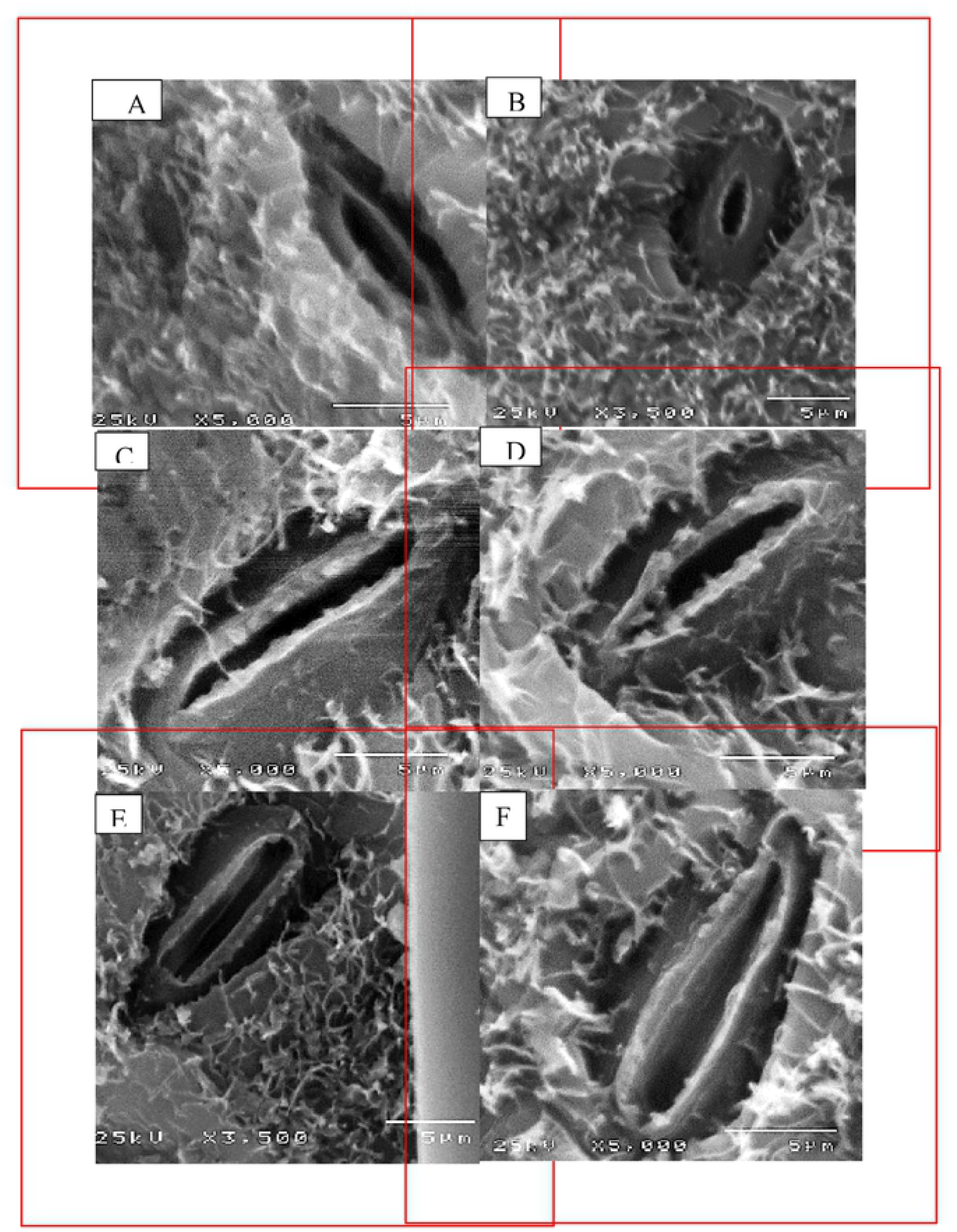
SEM photographs showing epiphytic PPFM on the stomata of Strawberry plant for all treatments. A: distilled water, B:10% methanol, C:30% methanol D: PPFM+10% methanol, E: PPFM+ 30% methanol, F: PPFM.

## Conclusion

Based on our findings, it can conclude that there were no significant differences between the two cultivars, except Florida c.v. was higher in fresh weight and early yield; however, Festival c.v. had higher total yield/plant and /fed.in the first season, anthocyanins, and ascorbic acid than Florida plants. When PPFM was sprayed, chlorophyll content, fresh weight, total yield /plant and /fed., TSS, anthocyanin, and ascorbic acid were all highest, whereas leaf area, dry matter percent, early yield, firmness, K, and carbohydrates were all highest when either PPFM or mixed with methanol 10% were treated. The best interaction was identified with Festival c.v. and PPFM treatment for total yield and most fruit quality features.

## Acknowledgment

The authors would like to thank the technical support from Horticultural Research Institute, Agricultural Research Center (ARC) and Faculty of women for Arts, Science and Education, Egypt

